# Revolutionizing male contraception: Personal lubricants as a novel way to deliver the Human Contraception Antibody

**DOI:** 10.1101/2025.08.14.670442

**Authors:** JM Doud, MT Geib, JA Politch, K Whaley, DJ Anderson, JG Marathe

## Abstract

**Study Question:** Can personal lubricants effectively deliver the sperm-agglutinating Human Contraception Antibody (HCA) to achieve on-demand male contraception?

**Summary Answer:** This study demonstrates that several water-based lubricants can effectively deliver bioactive HCA, and that a dimethicone-containing silicone lubricant can be modified into a stable emulsion suitable for antibody delivery.

**What Is Known Already:** The HCA-based vaginal film ZB-06 was shown to be safe and effective in a Phase I clinical trial for female contraception. Male contraceptive options remain limited. Using sexual lubricants as a delivery platform for HCA represents a novel and practical approach to male contraception.

**Study Design, Size, Duration:** We tested the stability of HCA for up to two years in a variety of commercially available sexual lubricants, and its delivery from a penile device using a simulated intercourse model.

**Participants/Materials, Setting, Methods:** Fourteen commercial lubricants were initially screened for impact on sperm motility and viability, miscibility with HCA solution, and preservation of HCA activity (sperm agglutination test). From these, three water-based and one silicone-based lubricant were selected for further study. The silicone-based product was engineered into a novel emulsion (KY-DE) by incorporating 6% w/w PEG-10 dimethicone and 0.005% v/v TWEEN-20 for HCA integration. Formulations were tested for HCA functional stability and contraceptive efficacy over time using kinetic sperm agglutination assays and a capillary tube sperm penetration test. A novel simulated intercourse model was developed using a 3D-printed penile device and a vaginal analog (Fleshlight®). HCA-lube was applied to the “penis”, and concentrations of HCA delivered to the “vagina” during intercourse were measured by ELISA and sperm agglutination assay. Safety evaluations were conducted using a vaginal tissue model (EpiVaginal, VEC-100-FT). Viability (MTT assay), tight junction integrity (TEER), and cytokine release were assessed after application of HCA-lube.

**Main Results and the Role of Chance:** All selected lubricants were nontoxic and did not affect HCA’s ability to agglutinate sperm. In the capillary tube sperm penetration assay, HCA-formulated lubricants significantly impaired sperm penetration; few motile sperm penetrated past a depth of 1 cm after 90 minutes in the HCA-lubricants vs >80_in most of the untreated lubricants. Importantly, the simulated intercourse assay confirmed successful delivery of HCA to the vagina. Water-based lubricants delivered an average of 24 ± 8.4 µg/mL of HCA, the commercial silicone lubricant 11 ± 5.2 µg/mL, and the KY-DE emulsion 27.4 ± 22.4 µg/mL. Futhermore, the formulations did not compromise tissue integrity or viability, or induce significant inflammatory responses.

**Large Scale Data:** Not applicable.

**Limitations, Reasons for Caution:** While these in vitro results support the feasibility of using lubricants as HCA delivery vehicles, translation into clinical application requires further validation and clinical trials.

**Wider Implications of the Findings:** This work presents a new direction in male contraceptive development, leveraging the widespread use of personal lubricants to introduce antibody-based, reversible, on-demand contraception. Engineering a silicone-based lubricant that retains HCA activity under varied conditions and ensures higher mucosal delivery could further advance this approach.

**Study Funding/Competing Interest(s):** This work was supported by a Sokol Grant from the Male Contraceptive Initiative, a Sexual Medicine Grant from Boston University Chobanian & Avedisian School of Medicine, and grant P50HD096957 from the National Institutes of Health. Kevin Whaley (ZabBio) intends to commercialize HCA for contraception. None of the other authors have competing interests.

## INTRODUCTION

Despite the availability of many modern contraceptive methods, approximately 50% of pregnancies worldwide are unintended^1^. Most unintended pregnancies occur in women that do not use modern contraception, or use it infrequently or improperly, and about 50% of unintended pregnancies result in abortion that can adversely affect maternal and child health^2,3^. Advances in contraceptive technology have predominantly focused on hormonal methods for women. Although men play a role in 100% of unintended pregnancies, male contraceptive options have remained largely unchanged^4^ and are limited to condoms, withdrawal, and vasectomy. Globally, condoms are utilized by approximately 21% of men, withdrawal by 5%, and vasectomy by 2%^5,6^. While condoms offer dual benefits of pregnancy prevention and protection against sexually transmitted infections (STIs), they have a typical user failure rate of around 13%^7^ and many couples discontinue their use due to perceived reductions in pleasure or intimacy^8^. Withdrawal is less effective, with failure rates nearing 20%, and vasectomy, although highly efficacious (>99.5%), requires a surgical procedure and is considered permanent, and thus is not widely adopted^7^.

Efforts to develop male hormonal contraceptives began in the 1930s^9^, with renewed interest emerging in the 1970s^10^. One approach is based on the administration of exogenous androgens which suppress luteinizing hormone (LH) and follicle-stimulating hormone (FSH) via the hypothalamic-pituitary-gonadal axis, and reversibly inhibit spermatogenesis^11^. Current hormonal candidates for male contraception include combinations of androgens with progestins, such as testosterone with Nestorone (NES+TES)^12^, and newer agents like dimethandrolone undecanoate (DMAU)^13^. Despite progressing into clinical trials, these methods have been associated with side effects such as mood disturbances, metabolic changes, inter-individual and racial variability in sperm suppression, and prolonged onset and recovery periods^11^. Among the non-hormonal male contraceptives, the leading candidates in preclinical development are YCT-529, a retinoic acid receptor antagonist, and soluble adenyl cyclase (sAC; ADCY10) inhibitor which is a small-molecule with efficacy demonstrated in the mouse model^14^.

In contrast to hormonal or small-molecule strategies, immunocontraception offers a novel, non-hormonal, and potentially on-demand approach to male contraception. We previously demonstrated the clinical feasibility of using a vaginal film (ZB-06) containing an anti-sperm monoclonal antibody (mAb), the Human Contraception Antibody (HCA), for contraception in women^15^. This first-in-class immunocontraceptive targets a glycan epitope on sperm^16^. HCA potently mediates sperm agglutination and immobilization, and also impedes sperm penetration through cervical mucus, thereby preventing fertilization^17^. The ZB-06 film was tested in a phase I clinical trial that entailed an exploratory post coital test. The film was safe and prevented the penetration of motile sperm into midcycle cervical mucus after intercourse^15^. Given its promising performance in female contraception, we sought to evaluate the potential of HCA as a male-directed contraceptive using personal lubricants as the delivery system.

Although various delivery systems for male contraception have been explored, including oral formulations, transdermal gels, injections, and implants^4,11^, topical delivery via personal lubricants remains an underexplored yet accessible and user-friendly modality. In the United States, personal lubricants are regulated as Class II medical devices by the Food and Drug Administration (FDA)^18^ and are broadly categorized into four classes: water-based, silicone-based, oil-based, and hybrid formulations^19^. Water-based lubricants, introduced commercially in the early 20th century^20^, remain the most widely used due to their condom compatibility and ease of use^21^. Silicone-based lubricants form a persistent, hydrophobic film ideal for prolonged activity^22^ and are commonly used for anal intercourse^23^. Hybrid lubricants combine properties of water- and silicone-based formulations, offering a balance of longevity and sensory appeal^24^.

The formulation of these products is critical for their performance and safety. Water-based lubricants are generally comprised of water-soluble polymers, buffers, emulsifiers, and preservatives, functioning as viscoelastic hydrogels that shear thin under stress^25^. Silicone-based lubricants primarily contain poly(dimethylsiloxane) (PDMS) and may be modified with dimethiconol, cyclomethicone, or tocopherols to tailor viscosity ^25^ and texture ^20^. These silicone formulations are typically bio-inert and do not perturb vaginal pH or osmolarity, factors linked to increased susceptibility to bacterial vaginosis and STIs^25^. Lubricants and their excipients can have varying effects on the mucosa. Prior research has demonstrated that hyperosmolar lubricants can result in mucosal damage and irritiation^26,27^ and some lubricants, like KY jelly, are toxic to lactobacilli, the primary microbial species in a healthy vaginal microbiome^25^. Several ingredients contained in the lubricant formulations have been associated with increased risk of infections; for example, polyquaternium-15 has been associated with epithelial damage and heightened HIV acquisition risk^26^, chlorhexidine increased susceptibility to chlamydia in a mouse model^28^, and propylene glycol was associated with increased HSV-2 acquisition in the mouse model^29^. Additionally, parabens can be absorbed systemically^30^ and have been linked to carcinogenesis, especially breast cancer^31^, emphasizing the need for careful selection of the type of lubricant and the excipients used. These findings underscore the importance of designing formulations that are not only effective but also compatible with the reproductive tract’s physiological and immunological milieu.

In this context, we present a proof-of-concept study evaluating the incorporation of the Human Contraception Antibody (HCA) into personal lubricants for topical application as a male-directed contraceptive. This strategy leverages the widespread use and acceptability of personal lubricants to deliver a non-hormonal, on-demand contraceptive that acts locally at the site of sperm deposition, offering a novel addition to the limited landscape of male contraceptive options.

## METHODS

### Semen samples

This study was approved by the Institutional Review Board at Boston University Medical Center (Human Subjects Protocols H-36843 and H-41454). All participants provided written informed consent prior to participation.

Semen samples were obtained from self-reported healthy men between the ages of 18 and 45, after 48 hours of abstinence. Semen sample collection and processing was done as previously described^17^. Motile sperm were obtained by passage of semen through 90% ISolate density gradient medium (FujiFilm Irvine Scientific; Santa Ana, CA, USA). Sperm concentration and motility were assessed on a Computer-Assisted Sperm Analysis system (CASA; CEROS II system with Human Motility II software, Hamilton Thorne, Beverly, MA, USA). The motile sperm pellet was resuspended to obtain a sperm concentration ranging from 30-40 x10^6^/mL in either Multipurpose Handling Medium^TM^ (MHM) (FujiFilm Irvine Scientific, Santa Ana, CA) for Washed Sperm (WS), or seminal plasma for Sperm in Seminal Plasma (SSP).

### The Human Contraception Antibody (HCA)

HCA, a human IgG1 monoclonal antibody (mAb), was produced in *Nicotiana benthamiana* using the variable sequence of H6-3C4 as previously described^17^. The antibody was provided for this study by Mapp Biopharmaceutical, Inc., San Diego, CA.

### Screening OTC lubricants for suitability as antibody delivery vehicles

A variety of personal lubricants was purchased over the counter (OTC). Water-based, silicone-based, and hybrid lubricants were assessed as part of this study and are listed in Table 1.

**Table 1:** Commercial lubricants advanced to additional testing.

| Lube | Type | Supports<br>Sperm<br>Viability | Supports<br>Sperm Motility | Miscibility with<br>HCA Solution | Supports HCA<br>Agglutination | Comments |
| --- | --- | --- | --- | --- | --- | --- |
| LL | Water-based | + | ++ | + | + | Best amongst water-based lubricants in screening |
| KY-C | Water-based | + | + | + | + | Largest market share |
| KY-D | Silicone-based | + | ++ | - | + | Best amongst silicone lubricants in screening |
| BD | Water-based | + | ++ | + | + | Marketed for preserving fertility |

Positive and negative groups were based on comparison of lubricant treatment groups with medium treated control. (+ Sperm viability = <10% increase in sperm death) (+ Sperm motility = >75% motility compared to medium and ++ = >90% motility). All water-based lubricants support antibody miscibility (+ indicates no visible separation of phases), but the silicone-based KY-D did not support antibody miscibility (- indicates visible separation of phases). All lubricants supported retention of antibody activity (+ indicates time comparable to medium for 100% agglutination in 100 µg/mL HCA kinetic agglutination assay).

Lubricants were diluted 1:4 or 1:16 in MHM by vigorously mixing and vortexing. Viscous lubricants were measured and dispensed with serial pipettes or syringes for accurate measurements. Three tests were used to evaluate the lubricants:

a. Miscibility with MHM: Lubricants diluted with MHM were visually assessed to determine miscibility with MHM.
b. Sperm viability: Washed sperm were exposed to MHM-diluted lubricants for 30 seconds and then tested using a LIVE/DEAD Sperm Viability Kit, (ThermoFisher Scientific, Waltham, MA, USA) per manufacturer’s protocol. Live vs dead sperm were differentiated and counted by fluorescence microscopy.
c. Sperm motility: The motility of WS diluted in lubricants was evaluated with the CASA and normalized to the motility of WS in MHM.
d. Sperm Agglutination: The kinetic sperm agglutination assay was performed as previously described^17^. Lubricants, diluted 1:16 in MHM, were mixed with HCA to achieve serial antibody dilutions ranging from 100 to 6.25 μg/ml.

### Development of HCA-containing Silicone Emulsions

Silicone lubricants are neither soluble nor miscible in aqueous solutions and require the addition of surfactants to prevent phase separation. Based on its low viscosity, single-ingredient composition of dimethicone, and good compatibility with the HCA in screening, KY True Feel Deluxe silicone lubricant was selected as the base for engineering a novel HCA-containing silicone emulsion, KY-DE. A surfactant was required to emulsify the antibody solution. Compared to ionic surfactants, non-ionic surfactants are safer for application to skin^32^ and mucus membranes^33^ and compatible with monoclonal antibodies, maintaining their structural integrity^34,35^. Three non-ionic surfactants, Cetyl PEG/PPG-10/1 Dimethicone (G90, Gransurf 90, Grant Industries; Elmwood Park, NJ, USA), Cetyl PEG/PPG-10/1 Dimethicone, Hexyl Laurate, Polyglyceryl-4 Isostearate (Gransurf W9, Grant Industries; Elmwood Park, NJ, USA), and PEG-10 Dimethicone (G67, Grant Industries; Elmwood Park, NJ, USA) were evaluated for their suitability for making HCA/lubricant emulsions. G67 was chosen as the lead surfactant based on observed room temperature stability (macroscopic) and the most homogeneous size and morphology under phase contrast imaging, compared to other surfactant formulations (Figure 4).

Reverse micelles were generated to encapsulate HCA and create a stable water-in-silicone (W/Si) emulsion. The aqueous phase consisted of HCA in 1X PBS with 0.005% v/v TWEEN-20^34^, while the silicone phase contained KY True Feel Deluxe and the nonionic surfactant PEG-10 Dimethicone (G67) at 6% w/w within the silicone phase. In 9 mm screw-thread glass vials (Fisher Scientific; Waltham, MA, USA), the water phase and silicone phase were combined at a 1:3.3 w/w ratio. A low-water fraction (>30% water) emulsion^36^ was desired to maintain high antibody concentration within the water phase, reduce viscosity, prevent phase inversion, and increase stability. This prototype ratio was derived empirically by observing macroscopic stability after mixing. Emulsification was performed using a VialMix (Lantheus Medical Imaging; Billerica, MA, USA) in two rounds of 15 seconds each, separated by a two-minute cooling period (Figure 1). The resulting emulsion were assessed for structural stability (phase separation) and functional stability (HCA activity).

**Figure 1:**
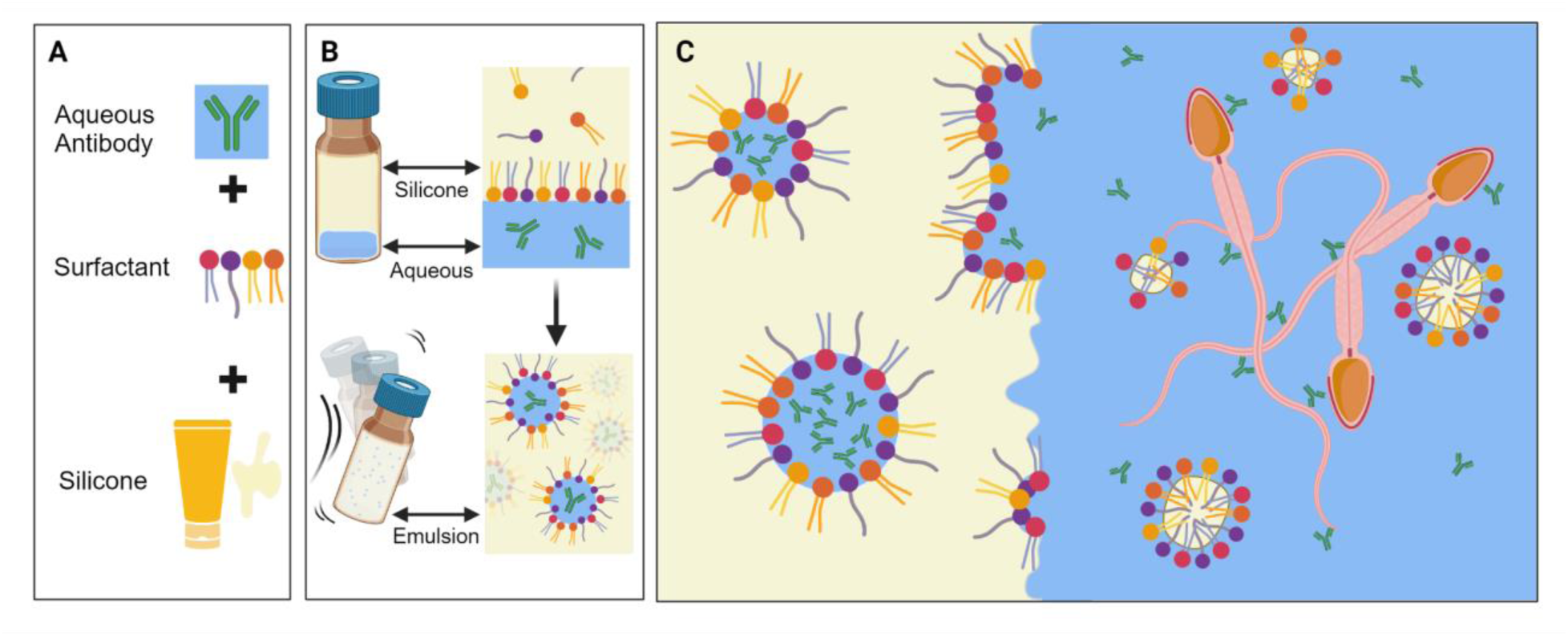
Engineering HCA-dimethicone emulsion with reverse micelles. A) To generate a stable emulsion, surfactants mediate the stability between aqueous and non-aqueous phases. B) Before mixing, aqueous and silicone phases are separate, with surfactant molecules adsorbing to the liquid interface. However, with mixing, these surfactant molecules encapsulate the dispersed aqueous antibody solution into reverse micelles within the silicone continuous phase. C) As bodily fluids, increased temperature, and shear forces are introduced to the emulsion, the reverse micelles will undergo phase inversion into traditional micelles containing silicone and releasing their aqueous payload into bodily fluids allowing HCA to agglutinate sperm.

### Evaluating material properties of HCA-containing Silicone Emulsions

#### a) Viscosity Verification

For selected OTC lubricants and HCA-dimethicone emulsions, the viscosity change over increasing shear rates at 37°C was analyzed using a DHR-2 Rheometer (40 mm, 2° cone and plate geometry, Peltier plate, TA Instruments; New Castle, DE, USA). Aliquots of emulsions and water-based lubricants were prepared with an HCA at a concentration of 1 mg/mL; emulsions were formulated as above, whereas aliquots of water-based lubricants received 46 µL of concentrated HCA or MHM (negative control). Before each test, the cone and plate were thoroughly washed with water and isopropanol. One mL of each sample was exposed to a constant shear stress of 100 Pa and sweeping shear rate from 1 to 100 per second after a 1-minute temperature equilibration soak at 37°C.

#### b) Structural Stability in HCA-containing Silicone Emulsions

To determine the optimal surfactant type and concentration, the homogeneity of the silicone emulsions was assessed as a thin film using phase contrast microscopy on an Axiovert S100 Inverted Microscope (ZEISS Group; Oberkochen, Germany). Surfactant concentrations of 2% to 32% w/w in the silicone phase was tested and formulated into 1 mg/mL HCA emulsions. A droplet of each prototype emulsion was placed onto a microscope slide (Frosted Microscope Slide 25 X 75 X 1mm; Fisher Scientific; Pittsburgh, PA, USA) and covered carefully with a square cover glass (Cover Glass 22 X 22 X 1 mm; Fisher Scientific; Pittsburgh, PA, USA) as to not induce shearing. The type and concentration of the lead surfactants were determined by observing homogeneity and average micelle size in the emulsion. Lead emulsions were selected for most comparatively monodisperse. For continuous monitoring of emulsion stability, silicone emulsions were examined visually (macroscopically) for signs of phase separation and flocculation. For long term monitoring, KY-DE was prepared at 1.6 mg/mL and stored at 4°C, room temperature (RT, 22-25°C), and 37°C. Emulsions were visually assessed at 0, 1, 3, and 30 days after initial preparation.

### Selection of Lubricants for additional testing

Three water-based [#LubeLife (LL), K-Y Jelly Classic (KY-C), and BabyDance (BD)] and one silicone-based [KY True Feel Deluxe (KY-D)] OTC lubricant were selected for additional testing. The determination to advance these lubricants was based on the results of initial screening (Table 2) and market-share/popularity among users.

### Evaluating HCA Stability in Lubricants and Silicone Emulsions using kinetic agglutination assay

To evaluate the temporal and temperature stability of HCA activity in lubricant formulations, time-course experiments were conducted using two vehicles: undiluted commercial lubricants and the silicone-based emulsion, KY-DE, described above. Stocks were prepared at an initial HCA concentration of 1.6 mg/mL and stored at three temperatures: 4°C, room temperature (RT), and 37°C. Aliquots were collected at designated time points—days 0, 1, 3, and 30 for both HCA-lubricants and KY-DE, with additional sampling at days 90 and 180 for HCA-lubricants to assess antibody function over time. Prior to analysis, all samples were diluted 1:16 in MHM to yield a final HCA concentration of 100 µg/mL. HCA-lubricant samples were assayed directly, whereas KY-DE samples required pre-processing due to optical interference from silicone reverse micelles. These emulsions were incubated at 37°C for 5 minutes, briefly vortexed, and centrifuged at 7000 RPM for 10 minutes to separate the aqueous and silicone phases. The lower aqueous phase was carefully aspirated and used for testing. Antibody activity was quantified using a kinetic sperm agglutination assay as previously described^17^.

### Sperm Penetration Assay

Flat capillary tubes (Borosilicate Capillary Glass Slide, 0.30x3.0 mm, 50 mm, Electron Microscopy Sciences, Hatfield, PA, USA) were marked at 1 cm intervals. Lubricants were diluted 1:4 in MHM, and HCA was added for a final concentration of 100 µg/mL. KY-DE was prepared at 1 mg/mL HCA and diluted 1:8 in MHM for a final antibody concentration of 125 µg/mL. The diluted HCA-containing lubricants were aspirated into the capillary tubes, and one end was sealed with parafilm or nail polish to prevent leakage. Sperm penetration into the capillary tube was assessed as previously described for cervical mucus penetration^37^.

### 3D construction of a Penile Device

A penile device was modeled in SOLIDWORKS® 2022 EDU using average dimensions for penile circumference of 12.23 cm and length of 14.15 cm^38^. It was designed to include features similar to the frenulum and glans to promote mixing. Additionally, this device was modeled with an internal cavity to house a 3cc BD Plastipak syringe, and a smaller cavity, modeled to resemble a urethra, into which a 21 x 3/4 gauge OD catheter with its needle removed (Butterfly® 21 X ¾ 12” Tubing; Abbott Laboratories, North Chicago, IL, USA) was fed through. This penile model was exported to UltiMaker Cura and 3D printed using an Ender 3 V2 FDM 3D printer. The device was printed in PLA using a 0.5 mm nozzle size at a standard 0.2 mm resolution. Both the .stl and Cura files are included in the supplemental materials.

### Assessing HCA delivery by Simulated Intercourse (SI)

An experimental setup was designed to simulate intercourse and assess delivery of HCA to the vaginal mucosa upon application of the HCA-dilute lubricant/ silicone emulsion to a penile device (Figure 2). Lubricants with an HCA concentration of 1 mg/mL were prepared and 0.75 mL was applied to the penile device. Methylcellulose was used to mimic vaginal secretions^39^. 4000 cP methylcellulose was prepared by adding methylcellulose (Sigma, Burlington, MA, USA) at 10 mg/mL in 1x EBSS (Thermo Fisher, Waltham, MA, USA). Methylcellulose was deposited 4 to 4.25 cm into the vaginal model (Ice Lady Fleshlight, Fleshlight, Austin, TX, USA). In KY-DE experiments, the vaginal model was lined with a female condom (Ormelle Female Condom, Cupid Limited, Nashik, India) for ease of sampling, and 300µL of methylcellulose^40^ was deposited directly into the condom. The lubricated penile device was thrust 10 times into the Fleshlight to a depth of 4.5 cm. Samples were taken from various locations along the vaginal canal with ophthalmic wicks (Merocel Eye Spear, Medtronic Xomed, Jacksonville, FL, USA). The samples were eluted from the swabs in 300 µL PBS and stored at 4°C overnight.

**Figure 2:**
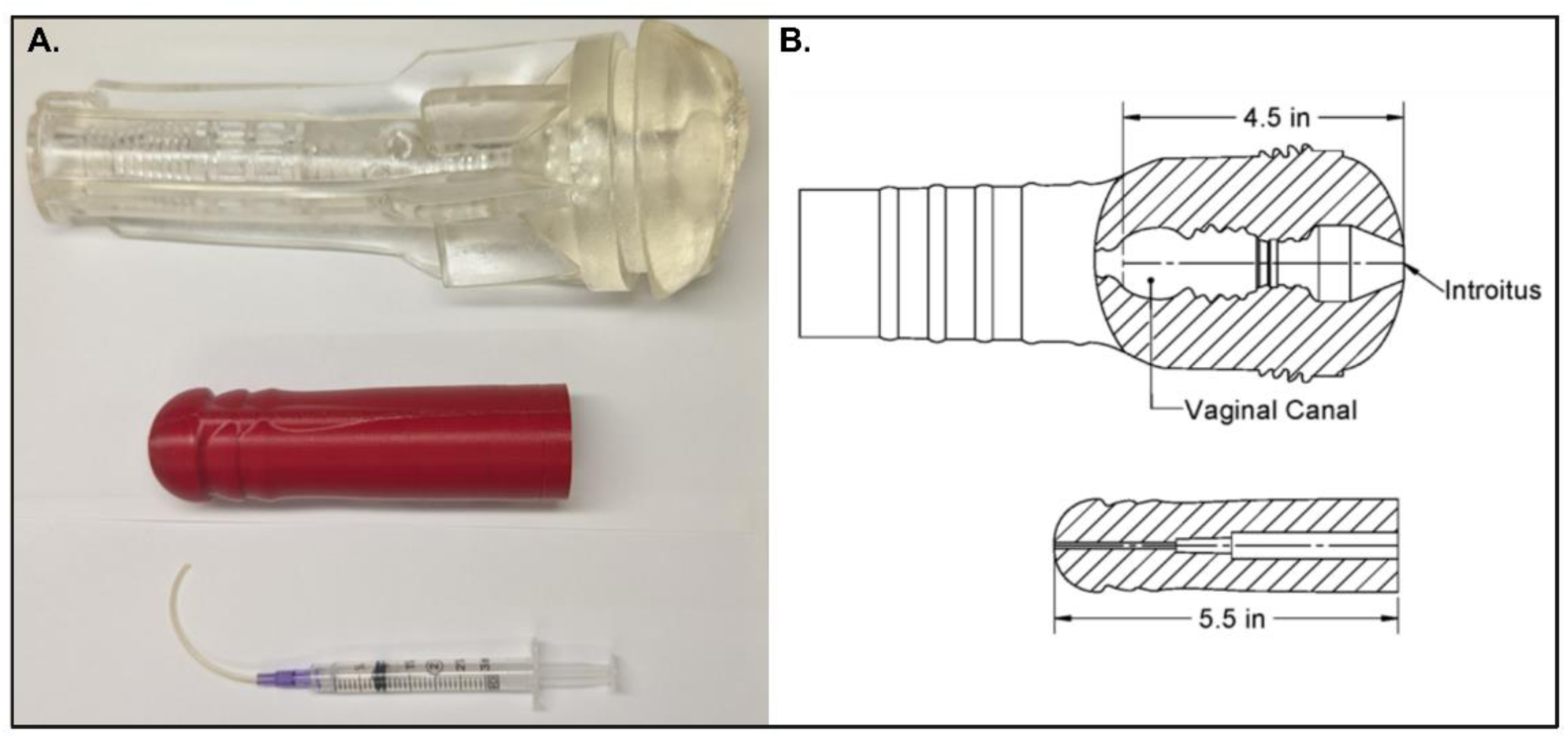
Simulated intercourse model incorporating a 3D-printed penile device and a vaginal canal analog (Fleshlight). A) Simulated intercourse experiments were designed using a Fleshlight (top), penile device (middle) and a tubing attached to syringe for injection of sperm (bottom). B) Schematic diagram of the length wise cross-section of Fleshlight (top) and 3D constructed penile device (bottom) with the dimensions representing physiological dimensions.

#### a) Measurement of HCA concentration by ELISA

HCA concentrations from the SI samples were measured with a modified in-house HCA ELISA^15^. Briefly, 1.5x10^5^ washed sperm in 50µl of MHM were added to each well of a clear polystyrene Corning^®^ ELISA 96-well microplate. The plate was incubated overnight, spun down at 300g, and the supernatant was discarded before the plate was allowed to air dry for 2 hours at 37°C. Eluted samples from the simulated intercourse test were added in duplicate alongside an HCA standard curve, and the remainder of the assay was performed as previously described^15^. The HRP-colorimetric signal and OD values were read at 450nm on a BioTek Synergy HTX multi-mode plate reader (Agilent, Santa Clara, CA, USA). HCA concentration was quantified by interpolating the standard curve (sigmoidal 4PL curve fit) (R2: 0.9985).

#### b) In vitro post coital test

1.2 mL of sperm in seminal plasma (SSP) was aspirated into a 3cc syringe, which was then attached to a catheter. The catheter was threaded through the “urethra” of the penile device and the syringe was inserted into the shaft. Any excess tubing length was cut off so the end of the catheter was flush with the tip of the penile device. SI experiments were performed as described above. The SSP was deposited in the vaginal canal after the initial 10 thrusts by depressing the syringe plunger. After five additional thrusts, the “post-coital” samples were taken from the cervix. Three µL of the sample was loaded on a sperm counting chambered CASA slide (DuraSlide CASA Non-Gridded Sperm Counting Chamber 20 µm Depth, Hamilton Thorne, Beverly, MA, USA). A total of 9 fields per sample were visualized at 20X magnification and were recorded with the CASA. Motile sperm were manually counted by a trained technician from the frames recorded on CASA to ensure accuracy as the automated program frequently counts agglutinated sperm as motile.

### Evaluating the effects of HCA-lubricant on epithelial tissue

Full-thickness EpiVaginal^TM^ (VEC-100-FT) (MatTek, Ashland, MA, USA) were used as vaginal and foreskin 3D organotypic model to perform irritation studies. Tissues were transferred to 24-well plates with 500 µL of the supplied MatTek medium on the basal side, and 50 µL medium on the apical side. Tissues were then equilibrated overnight in an incubator at 37°C. The following morning, 50 µL of HCA-lubricants diluted 1:4 in MHM were added to the apical side of the tissues and incubated for 4 hours at 37°C. Lubricants were gently aspirated and removed, tissues were washed MatTek tissue medium and then 50 µL of medium was added, and tissues were incubated overnight. Apical and basal supernatants were collected after 24 hours and frozen at -80°C for future cytokine analysis.

#### a) TEER measurement

Changes to epithelial barrier integrity were evaluated using trans-epithelial electrical resistance (TEER) measurements. All tissues were evaluated at baseline and 24 hours after lubricant application with the EVOM3 (WPI, Sarasota, FL, USA) per manufacturer instructions. TEER measurements were normalized to untreated control tissues

#### b) MTT Assay

MTT [3-(4,5-dimethylthiazol-2-yl)-2,5-diphenyltetrazolium bromide] (Sigma, St. Louis, MO, USA) assay was performed per manufacturer’s instructions. Concentration of formazan formed by metabolically active cells was measured with optical density (OD) at 570nm using Biotek Synergy HTX plate reader (Agilent, Santa Clara, CA, USA). Change in tissue viability was determined by comparing OD of treated tissues to that of untreated tissues.

#### c) Cytokine Analysis

Inflammatory cytokine levels in apical and basal supernatants were analyzed with a 13-plex ProcartaPlex kit (Life Technologies, Grand Island, NY, USA) according to manufacturer’s instructions. The analytes included IL-1 alpha, IL-1 beta, IL-1RA, IL-6, IL-7, IL-8 (CXCL8), IP-10 (CXCL10), MIP-1 alpha (CCL3), MIP-1 beta (CCL4), MIP-3 alpha (CCL20), RANTES (CCL5), and TNF alpha and were assayed by Luminex (Bio-Plex MAGPIX, Bio-Rad Laboratories, Hercules, California, USA).Cytokine levels from treated tissues were compared to those of untreated (medium) controls.

### Statistical Analysis

All experiments were done with at least 2 replicates and repeated at least 3 times. For assays using sperm, three individual donors were used for each experiment. GraphPad Prism (Version 10.4; GraphPad Software Inc.; San Diego, CA, USA) and JMP Pro (Version 18.0.2; SAS Institute, Cary, NC, USA) were used for statistical analysis and graphing of data. Data were tested for normality by the Shapiro-Wilk test and were log transformed prior if determined not to be normally distributed. Data were then subjected to analysis of variance (ANOVA) or mixed effects analysis. A significant omnibus test was followed by post hoc Tukey’s or Šídák’s multiple comparison tests. For analysis of cytokines, data from lubricants were expressed as percent of respective medium controls and then analyzed by Kruskal-Wallis ANOVA. A significant ANOVA was followed by Dunn’s post hoc multiple comparison test. Differences were considered to be statistically significant when p < 0.05.

## RESULTS

### 1. Suitability of commercial lubricants for use as a vehicle for antibody delivery

We evaluated 14 commercial lubricants (Table 1 and Supplemental Table 1). As expected, water based (6/6) and hybrid lubricants (2/3) were miscible with water-based HCA solution and silicone based lubricants (4/5) were not. We noted that Wet-Water-based lubricant was viscous and not miscible with HCA solution while the Swiss Navy Silicone was miscible. Sperm viability was variable in water-based lubricants and 4 out of 7 lubricants resulted in sperm immotility and death. The majority of the silicone-based lubricants (4/5) preserved sperm viability. HCA maintained its activity in all lubricants except the hybrid lubricants in which neither the sperm viability nor the agglutination could be accurately assessed due to generation of microbubbles limiting visibility under the microscope. Based on initial screening, 4 representative lubricants that support sperm viability, motility, and HCA agglutination were selected (Table 1). These include 3 water-based lubricants: #LubeLife (LL), KY Classic (KY-C), BabyDance (BD) and 1 silicone-based lubricant: KY Deluxe (KY-D).

### 2. HCA Silicone Emulsion Characteristics

#### a) Verification of viscous properties

Studies have estimated shear rates during intercourse to range from as low as 0.1 s⁻¹, driven by capillary forces, up to values exceeding 100 s⁻¹ during coitus ^41-43^. To establish appropriate shear response criteria for a novel silicone emulsion lubricant (KY-DE), a representative shear rate range of 1–100 s⁻¹ was selected. The shear response of all lead lubricants and KY-DE was evaluated using a DHR-2 Rheometer, measuring viscosity as a function of increasing shear rate at 37°C (Figure 3). All water-based lubricants (LL, BD, KY-C) in the absence of HCA exhibited shear-thinning behavior, characterized by high viscosity at low shear rates and lower viscosity at high shear rates. Their viscosity decreased from a range of 346–23,025 mPa·s to 157–1,233 mPa·s. In contrast, the silicone-based lubricant KY-D behaved as a near-Newtonian fluid, maintaining a consistent viscosity with a mean of 38.5 ± 2.4 mPa·s across the tested shear rates. KY-DE demonstrated shear-thinning behavior with a more pronounced decrease in viscosity, from 1,560 to 88 mPa·s. At low shear rates, KY-DE resembled a water-based lubricant in viscosity but progressively thinned with increasing shear, approaching the viscosity of its base component, KY-D. A small, non-significant reduction in average viscosity was observed between formulations without HCA (Figure 3A) and those containing HCA (Figure 3B), attributable to dilution with the aqueous HCA solution. However, no major differences in rheological behavior were noted following HCA incorporation.

**Figure 3:**
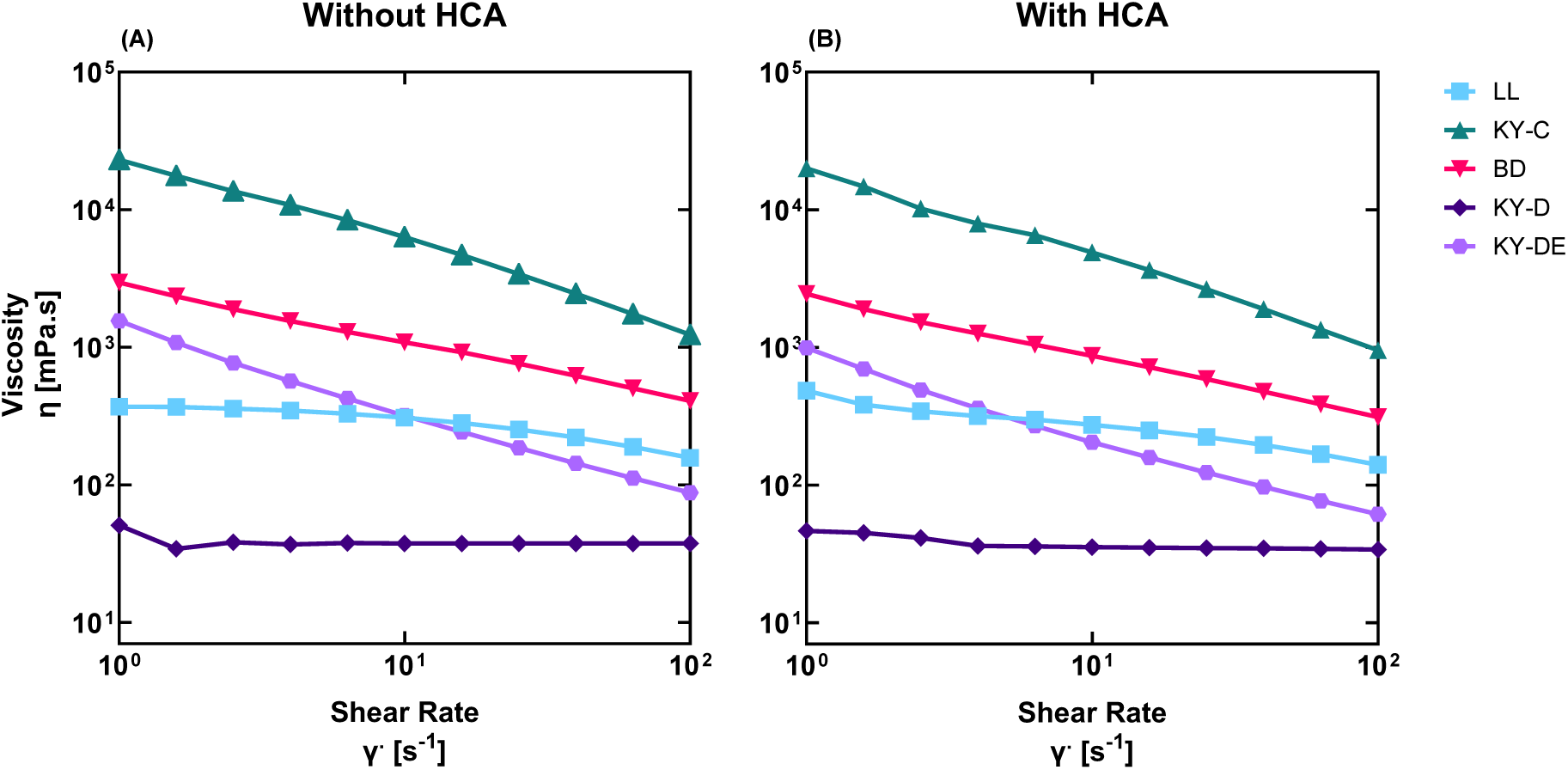
Evaluating viscous properties of lubricants. Shear thinning properties in silicone, silicone-emulsion and water-based lubricants. Shear flow response was measured in LL, KY-C, and BD, KY-D, and KY-DE using a DHR-2 Rheometer. Testing was conducted at 37 °C while exposed to a constant shear stress of 100 Pa and sweeping shear rate from 1 to 100 per second. Under increasing shear rates, LL, BD, and KY-C (water-based lubricants) behave as non-Newtonian shear-thinning fluids. KY-DE (W/Si emulsion) also shear thins while KY-D (silicone-based lubricant) behaves as an ideal Newtonian fluid. All lubricants behaved similarly, both A) without HCA and B) with 1mg/mL HCA.

#### b) Emulsion macrostructure and stability

Under 40X phase contrast observation, G67 and G90 surfactants at 2-6% w/w within the silicone phase produced the most monodisperse distribution of micelles with the smallest average size. G90 4%, G67 4%, and G67 6% were determined as lead candidates. Figure 4 shows representative stable and unstable emulsions in Panel I and Panel II, respectively.

**Figure 4:**
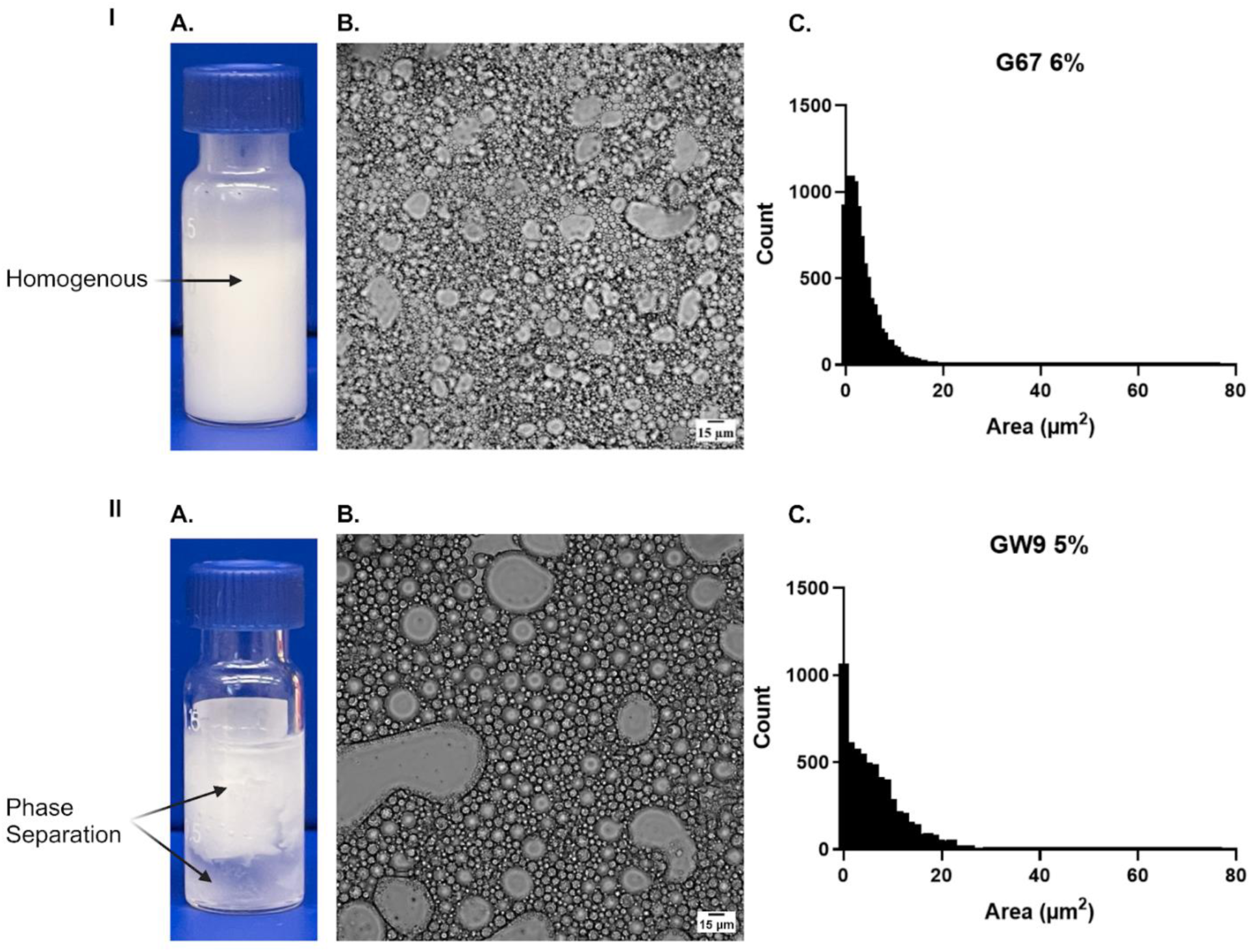
Visual and microscopic assessment of engineered emulsion stability. Representative data are shown for evaluation of emulsion stability. Panel I represents stable emulsion and Panel II shows phase separation of unstable emulsion. IA and IIA show the visual difference between stable and unstable emulsion. IB and IIB represent the phase contrast images (40X) taken at 24 hours with IIB demonstrating larger clear spaces where the water phase has separated compared to IB. In panel C, the size (μm^2^) distributions of micelles correspond to images in panel B, analyzed via ImageJ particle analyzer.

Emulsions were binarily categorized as stable or unstable (phase separation, aggregation, etc.) by their macroscopic properties by visual examination; emulsions made with G67 6% w/w were stable at 4°C and 25°C for at least 30 days. Full phase separation was observed after just 3 days at 37°C. All other emulsion permutations became unstable at 37°C and lower temperatures within 30 days.

### 3. Stability and Activity of HCA in Lubricants

#### a) Effect of temperature

Time-course experiments evaluating HCA stability across different temperatures revealed that HCA activity remained stable at lower temperatures (4°C) for up to 2 years. Within each lubricant type (LL, KY-C, BD and KY-D) as well as medium control, sperm agglutination rates did not differ from baseline (Day 0) until Day 720 at which point all lubricants and control were significantly slower than baseline (p<0.0001). At each storage duration time point, LL, KY-C and BD (but not KY-D) exhibited slower sperm agglutination rates compared to medium control (Figure 5A). KY-DE has only been assessed up to 90 days at the current time, but has shown excellent agglutination at 4°C.

**Figure 5:**
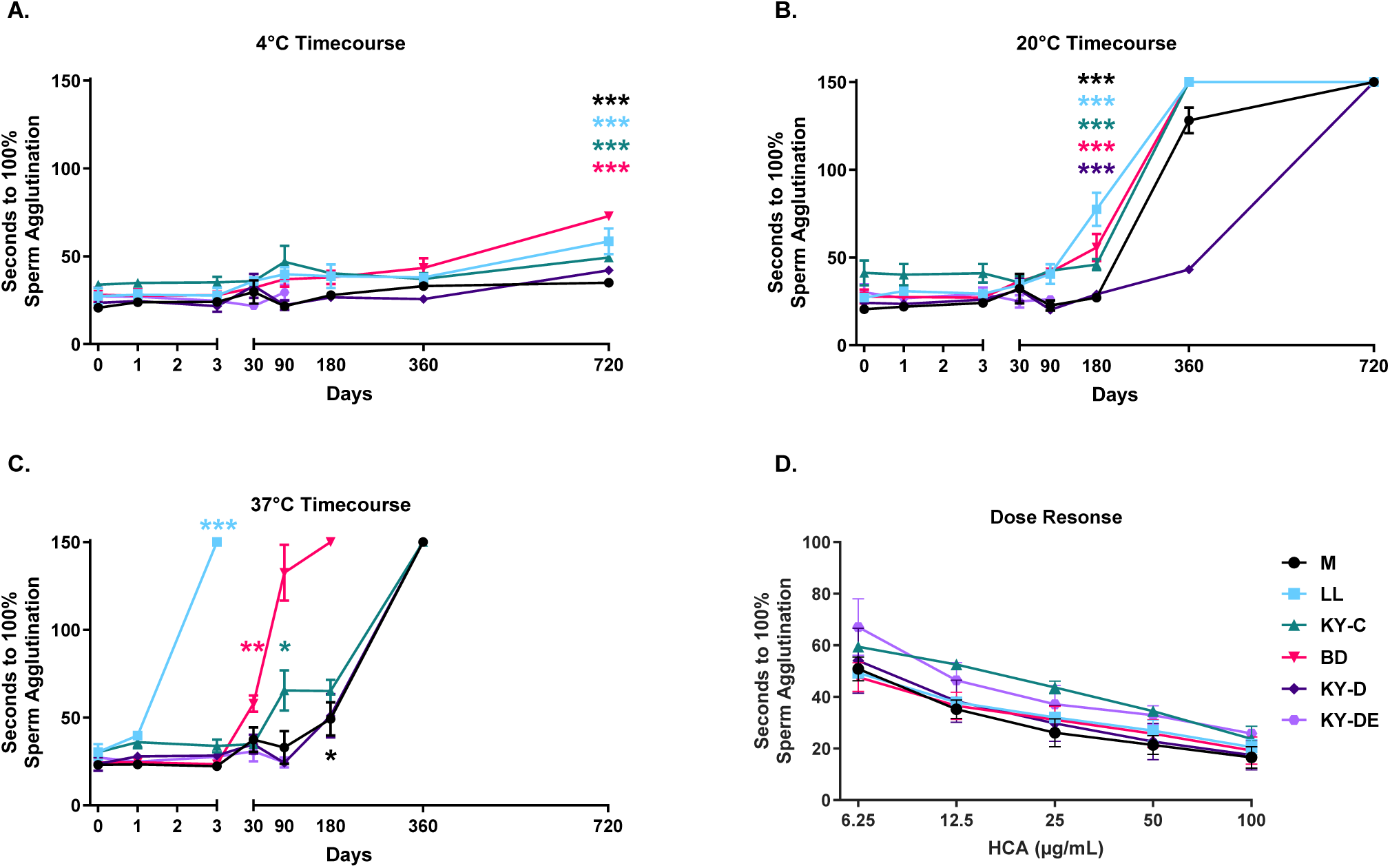
Stability and dose-response of HCA in lubricants. HCA activity was measured by kinetic sperm agglutination assay after storage of HCA-lubricants/emulsion at three different temperatures: 4°C, 20°C, and 37°C. The HCA-lubricant and emulsion stocks were diluted to HCA concentration of 100 µg/mL and were tested for time to 100% sperm agglutination. A) At 4°C, HCA activity was stable up to 360 days but declined somewhat at 720 days across all lubricant formulations compared to baseline (Day 0). B) At 20°C, a significant decline in HCA activity was observed in M, LL, KY-C and BD at 180 days, with complete loss of activity by 360 days in LL, KY-C, and BD. KY-D retained activity up to 360 days. C) At 37°C, HCA activity declined rapidly, with significant loss in agglutination as early as Day 3 for LL, and by Day 30 for most other lubricants. D) Dose-response of HCA activity was assessed by agglutination assay using serial dilutions of HCA (100 to 6.25 µg/mL) in 6.25% lubricant. Time to 100% agglutination increased significantly at lower HCA concentrations (p=0.001), but no significant differences were observed between lubricants (p=0.16). Data are shown as mean ± SEM and represent 3 independent experiments. Statistical significance: *0.002, **0.0002, ***0.0001

At room temperature (20°C) (Figure 5B), all lubricants (LL, KY-C, BD and KY-D) and medium control exhibited a significant reduction in HCA sperm agglutination activity at 6 months compared to baseline (p<0.0001). LL, KY-C, and BD completely lost agglutination activity by 1 year, while the HCA-medium control showed a notable increase in agglutination time, rising from an average of 20.47 seconds to 128.19 seconds. The silicone-based KY-D formulation with HCA exhibited complete loss of activity at 2 years. Similar to 4°C above, LL, KY-C and BD (but not KY-D) exhibited slower sperm agglutination rates compared to the medium control at each storage time point tested (i.e., up to 360 days). KY-DE has only been assessed up to 90 days at the current time but again has shown excellent agglutination activity at 20°C.

At elevated storage temperature (37°C) the loss of HCA sperm agglutination activity was accelerated in both HCA-lubricants and KY-DE formulations (Figure 5C). Significant differences in activity between lubricant types emerged as early as day 3 for LL and persisted over the full time course. Notably, LL demonstrated the most rapid loss of activity, with a complete loss observed by day 3, distinguishing it significantly from all other lubricants and the medium control. Though KY-C, BD, KY-D and KY-DE showed 100% agglutination at 3 months, only the KY-D (mean: 24.19 sec) and KY-DE (mean: 24.42 sec) had agglutination that remained similar to baseline (mean 23.59 sec). 100% agglutination with HCA in KY-C at 3 months was at 65 sec compared to 30.22 seconds at baseline while that in BD at 3 months was 132.56 sec compared to 24.22 sec at baseline on average. No HCA activity was observed at 6 months for BD and at 1 year for KY-C and KY-D. KY-DE has only been assessed up to 90 days at the current time but again has shown substantial agglutination activity at 37°C thus far.

#### b) Dose response

In a Kinetic Agglutination HCA titration assay, analyzed by 2-way ANOVA (Lubricant, HCA concentration), a significant main effect was found for increased HCA concentration (6.25, 12.5, 25, 50 and 100 µg/mL) (p=0.001) but no difference was detected between lubricants overall (p=0.16). We observed 100% sperm agglutination times, with all 100 µg/mL HCA lubricant solutions at 23-29 sec (Figure 5D).

#### c) Capillary Sperm Penetration Test

A modified sperm penetration test demonstrated that after 90 minutes, most of the lubricants containing 100 µg/mL HCA had ≤1 PMS/hpf at all distances; exceptions were M and LL, which had 2.1 and 1 PMS/hpf, respectively, at the 1cm distance. All values for the HCA-lubricants were below the contraceptive threshold of <5 PMS/hpf, whereas most of the untreated lubes had >5 PMS/hpf at all distances (Figure 6). Three-factor repeated measures ANOVA indicated significant main effects for distance (p<0.0001), HCA presence in lubricant/medium (p<0.0001) and for their interaction (p<0.0001) but not for lube type (p=0.10).

**Figure 6:**
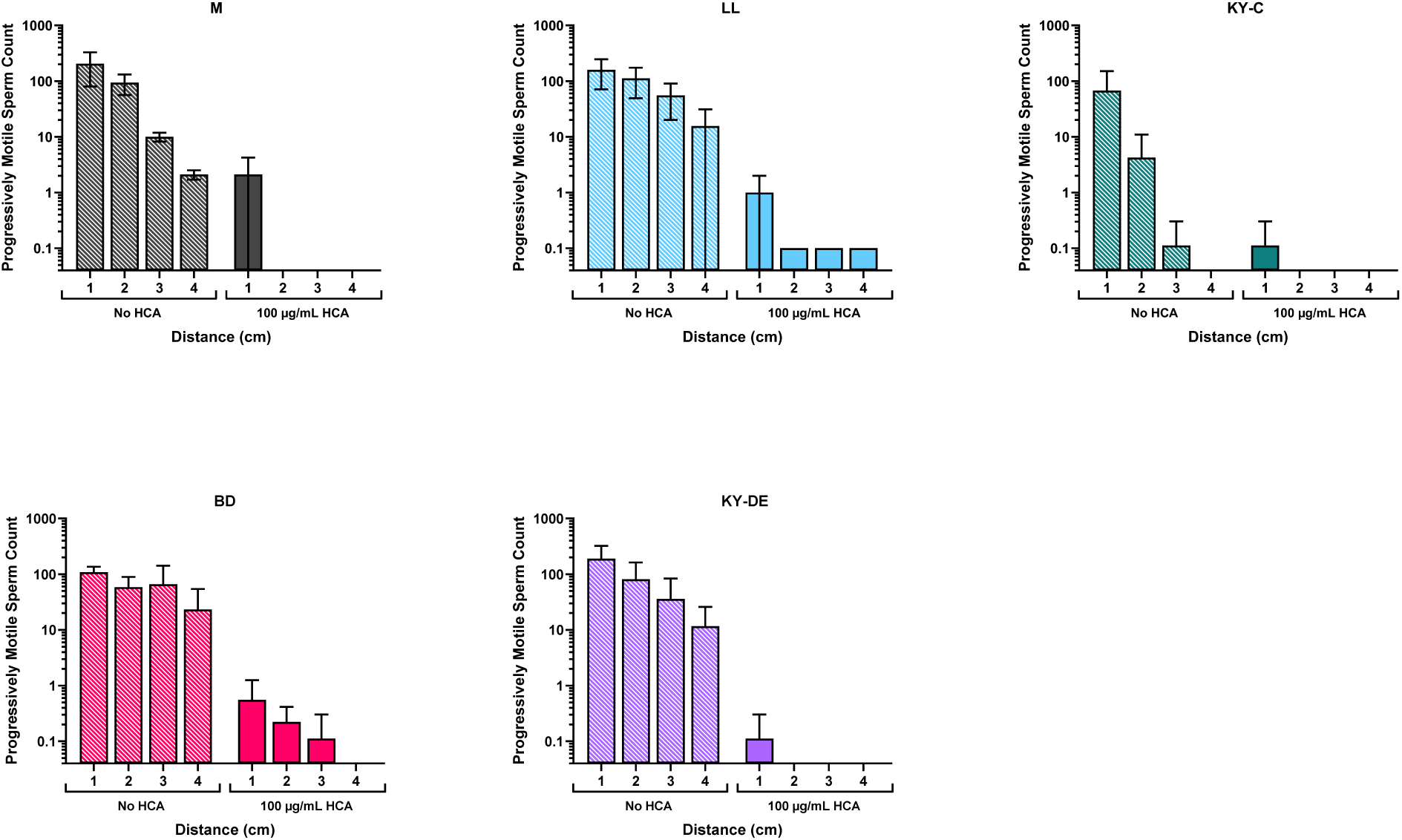
Sperm penetration through lubricants with and without HCA. A modified capillary tube sperm penetration assay was used to assess the ability of sperm to penetrate through lubricants with and without HCA. After 90 minutes, few sperm were observed to have penetrated through lubricants containing HCA compared to lubricants without HCA. Three-factor repeated measures ANOVA indicated significant main effects for distance (p<0.0001), HCA presence in lubricant/medium (p<0.0001) and for their interaction (p<0.0001) but not for lube type (p=0.10). Data are shown as mean ± SEM and represent 3 independent experiments.

### 4. Delivery and activity of HCA in HCA-lubricants and silicone emulsions in simulated intercourse experiments

#### a) Concentration of HCA delivered to the vagina

We observed that the concentration of HCA delivered to the Fleshlight’s vaginal canal by HCA-lubricant applied to the penis varied with the type of lubricant. The HCA concentrations (Mean±SEM) in samples from the introitus (I) vs vaginal canal (VC) were: LL 7.0±3.0 vs 11.2±3.4 µg/mL, KY-C 4.3±0.8 vs 5.7±1.0 µg/mL, BD 30.7±11.4 vs 31.4±4.9 µg/mL, KY-D 25.1±6.5 vs 40.0±10.6 µg/mL and KY-DE 43.7±6.6 vs 27.7±21.3 µg/mL. HCA concentrations measured by direct ELISA following simulated intercourse showed detectable concentrations of HCA in all lubricants. HCA concentrations in vaginal samples after delivery with LL and KY-C did not differ from one another but were both significantly lower than BD (p=0.019, 0.002), KY-D (p=0.016, 0.002) and KY-DE (p=0.049, 0.006), respectively. HCA concentrations did not differ significantly between the introitus and vaginal canal for any of the lubricants (p=0.76) following simulated intercourse (Figure 7A).

**Figure 7:**
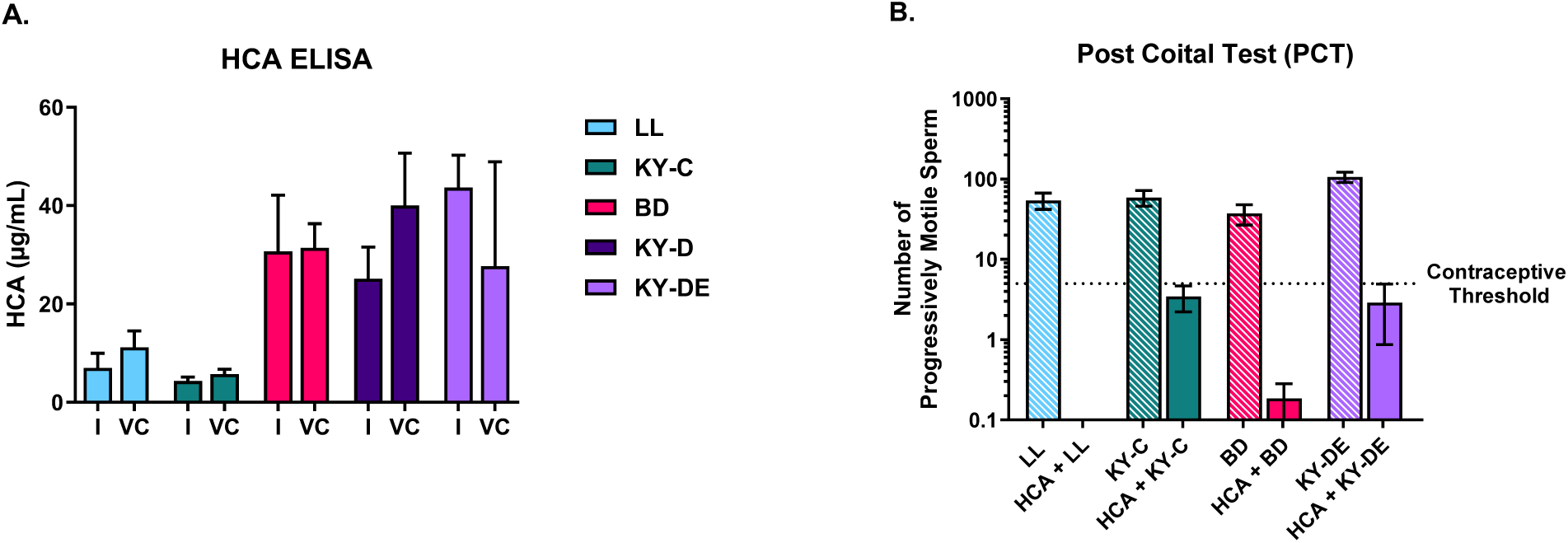
HCA delivery and activity in simulated intercourse experiments. A) HCA concentrations measured by direct ELISA (I= introitus and VC= vaginal canal) showed measurable concentrations of HCA in all lubricants. HCA concentrations in paired LL and KY-C samples did not differ from one another but were both significantly lower than BD samples (p=0.019, 0.002), KY-D (p=0.016, 0.002) and KY-DE (p=0.049, 0.006), respectively. HCA concentrations did not differ significantly at the introitus compared to the vaginal canal in paired samples for any of the lubricants (p=0.76) following simulated intercourse. B) PCT demonstrated significant decrease of progressively motile sperm (PMS) below the contraceptive threshold of <5PMS/ hpf in all lubricants containing HCA compared to lubricants without HCA. Data are presented as mean ± SEM and are representative of 3 individual experiments.

#### b) In vitro postcoital test (PCT) following simulated intercourse

Progressively motile sperm (PMS) were calculated per high power field (hpf) in vaginal samples after simulated intercourse (Figure 7B) (“in vitro PCT”). HCA-lubricants effectively inhibited PMS with mean counts below the contraceptive threshold of 5 PMS/hpf. KY-C + HCA resulted in 3.44 PMS/hpf, KY-DE + HCA resulted in 2.89 PMS/hpf, and LL and KY-D had ≤ 0.2 PMS/hpf. All Lubricants without HCA resulted in mean PMS/hpf exceeding the contraceptive threshold, with counts ranging from 37.26-121.33. The PMS/hpf of each HCA-lubricant condition was significantly lower than that of their respective lubricants alone: LL (p<0.005), KY-DE (p<0.0001), KY-C (p<0.005) and BD (p <0.05).

### 5. Tissue irritation studies

a. TEER and MTT assays were performed on lubricant-treated EpiVaginal tissue (Figure 8A, B) to evaluate epithelial integrity and tissue viability respectively. At 24 hours, there were no significant differences in lubricant-treated EpiVaginal tissues when compared with medium control.
b. We measured cytokine levels from apical and basal supernatants collected 24 hours after treatment of EpiVaginal tissues with the different lubricants. There were no significant increases in proinflammatory cytokine levels when compared with medium control (Figures 8C and D).

## Discussion

This study presents the first comprehensive evaluation of commercial personal lubricants as potential vehicles for the delivery of a contraceptive, HCA, in the context of on-demand male contraception. Our findings demonstrate that select commercially available water-based and silicone-based lubricants can support HCA activity, preserve sperm viability, and enable effective antibody delivery, suggesting a promising avenue for novel, non-hormonal contraceptive strategies.

**Figure 8:**
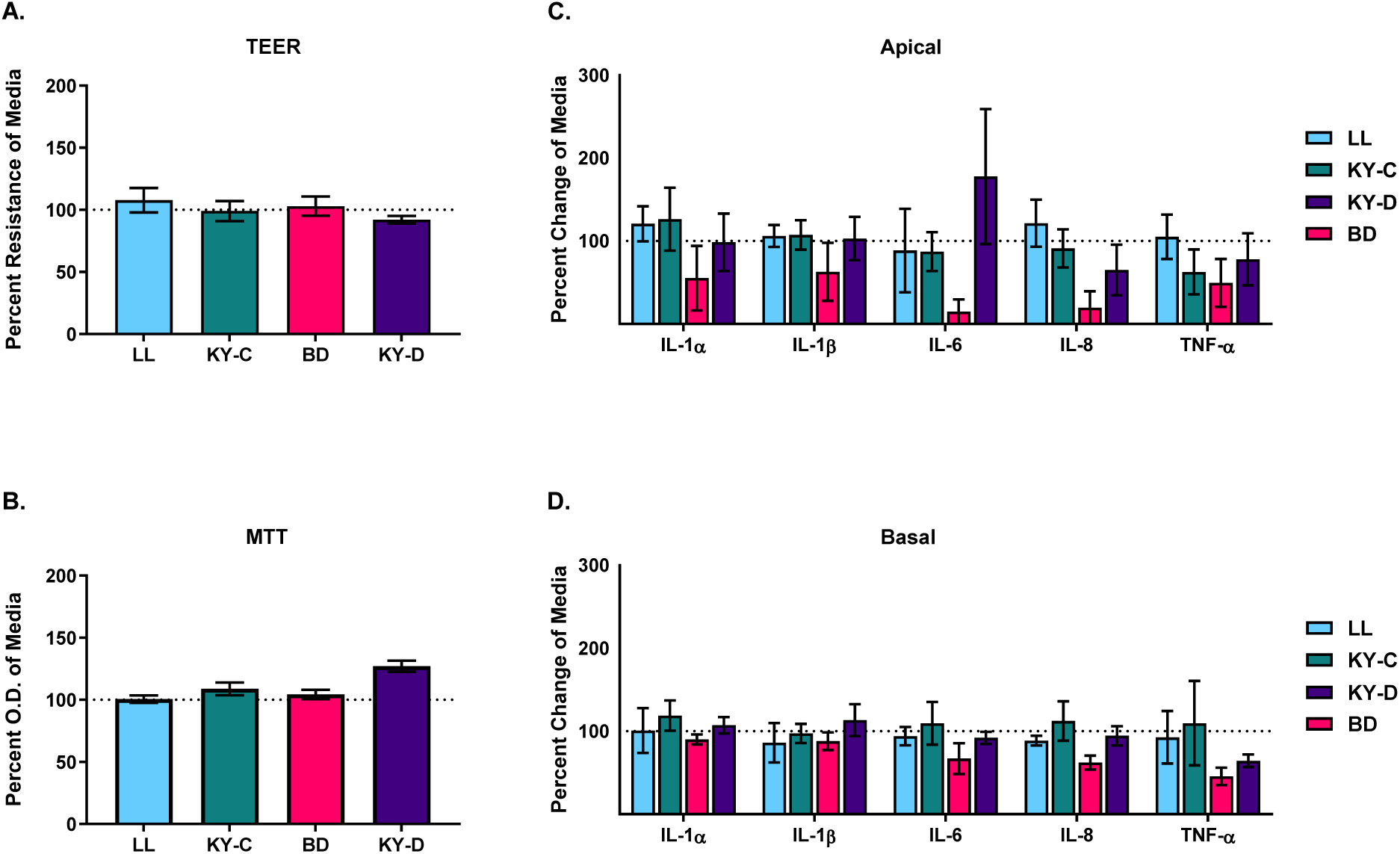
Tissue irritation tests. A) There was no difference in tissue integrity assessed by TEER measurements before lubricant exposure, and at 24 hours after a 4-hour lubricant exposure when compared to medium control. B) MTT assay showed comparable tissue viability across different lubricants. Measurements of proinflammatory cytokines showed no significant increases in concentration in apical (C) or basal (D) samples upon treatment of EpiVaginal tissues with lubricants as compared to medium control.

The initial screening of 14 lubricants revealed important insights into compatibility and functionality. While all water-based lubricants were miscible with aqueous HCA solution, only one silicone-based lubricant (Swiss Navy Silicone) showed unexpected miscibility. Conversely, water-based lubricants varied significantly in their ability to preserve sperm viability, with four of seven formulations causing complete sperm immobilization and death. These observations are consistent with prior literature indicating that osmolality and pH of lubricants can markedly influence sperm function^44,45^. Silicone-based lubricants demonstrated superior compatibility with sperm, aligning with their typical iso-osmolar and inert formulations. Recognizing the hydrophobic nature of silicone lubricants, we engineered a stable emulsion (KY-DE) incorporating 6% w/w PEG-10-Dimethicone to encapsulate aqueous HCA. Rheological characterization confirmed that KY-DE exhibits shear-thinning behavior, mimicking the dynamic changes in viscosity in water-based lubricants observed during coitus^41,42^. This behavior may facilitate the lubricant’s spreadability and comfort during use, while still enabling antibody delivery. Emulsion stability at 4°C and 25°C was achieved with G67 6% formulations, although storage at 37°C compromised structural integrity, highlighting the need for further formulation optimization for tropical and high-temperature environments.

Preserving the functional integrity of HCA over time is critical for product viability. Our time-course experiments demonstrated that HCA retained full activity for up to two years at 4°C, regardless of the lubricant type. However, room temperature storage led to significant degradation, especially in water-based lubricants, with complete activity loss within a year. Elevated storage temperatures (37°C) further accelerated this degradation. These findings align with known stability limitations of proteins in non-ideal storage conditions^46^, suggesting the necessity of cold-chain storage or further stabilization strategies.

Using a novel intercourse simulation model, we confirmed successful delivery of HCA from penile application into the vaginal canal. Notably, all lubricants with HCA achieved delivery concentrations above 6.25 µg/mL, exceeding previously determined thresholds for effective sperm agglutination. The engineered emulsion (KY-DE) achieved the highest delivery concentration but was not significantly different from commercial formulations, indicating that simple aqueous formulations may suffice for effective delivery but also that silicone emulsions may offer choice to suit user preferences. All HCA-lubricant combinations significantly reduced progressively motile sperm (PMS) counts in post-coital testing to below the contraceptive threshold (<5 PMS/hpf), a critical functional benchmark for contraception.

Biocompatibility is a key consideration for any topical vaginal or penile product. Safety studies were conducted *in vitro* using MatTek EpiVaginal (human vaginal tissue) models, which meet the European Centre for the Validation of Alternative Methods (ECVAM) and the Organization for Economic Co-operation and Development (OECD) guidelines for skin irritation, corrosion and permeability testing and are accepted as an alternative to animal testing^47^. Our safety assays, including TEER and MTT measurements, demonstrated that neither HCA nor the base lubricants caused epithelial damage or viability loss in EpiVaginal tissue models. Additionally, cytokine profiling showed no induction of a pro-inflammatory response, underscoring the non-irritating nature of both commercial and engineered formulations. These findings support the suitability of HCA-containing lubricants for mucosal application, consistent with existing data on HCA safety in prior vaginal film trials^15^.

## Limitations and Future Directions

While the in vitro findings are compelling, study limitations must be acknowledged. First, the use of simulated intercourse with synthetic material models, while innovative, cannot fully recapitulate in vivo dynamics of lubricant distribution, mucosal interaction, and antibody function in mucosal tissues. Second, the stability of HCA in emulsions requires further optimization to ensure consistent activity in diverse climatic settings. Third, while our in vitro post-coital testing strongly supports contraceptive potential, future lubricant engineering studies may be necessary to improve lubricant adherence to penile skin and delivery of product to the vagina. Additionally, in vivo human trials will be essential to validate efficacy, safety and acceptability.

## Implications for Male Contraception Development

Current male contraceptive options remain limited, and interest in non-hormonal, user-controlled methods continues to rise. Our study introduces a paradigm shift by leveraging a growing consumer product—personal lubricants—as a discreet and acceptable delivery platform for antibody-based male contraception. This approach aligns with recent trends in multipurpose prevention technologies (MPTs), combining ease of use with effective contraception. Future work will focus on optimizing HCA-lubricant formulations for temperature stability, increasing mucosal delivery efficiency, and conducting early-phase clinical trials. HCA may be combined with mAbs directed against STIs such as HIV-1 and HSV-2 to produce contraceptive MPTs.

