## Supplemental Table 1 for "Revolutionizing male contraception: Personal lubricants as a novel way to deliver the Human Contraception Antibody"

**Supplement Table 1: List and composition of over the counter sexual lubricants used in this study**

| Name | Base | Key Ingredients |
|---|---|---|
| #LubeLife | Water | Water, Propanediol, Gluconolactone, Hydroxyethylcellulose, Sodium Bensoate, Citric Acid |
| Astroglide Glycerin & Paraben Free | Water | Purified Water, Butylene Glycol, Propylene Glycol, Xylitol, Polyquaternium 15 |
| Astroglide Liquid | Water | Purified Water, Glycerin, Propylene Glycol, Xylitol, Polyquaternium 15 |
| His & Hers Arousing and Tingling Personal Lubricant | Water | Dimethicone, Cyclopentasiloxane, Dimethiconol, Phenyl Trimethicone, Vanillyl Butyl Ether, Mentha Piperita (Peppermint) Extract |
| K-Y Jelly Classic | Water | Water, Glycerin, Hydroxyethylcellulose, Gluconolactone, Methylparaben, Sodium Hydroxide, Chlorhexidine Digluconate |
| K-Y True Feel Deluxe | Silicone | Dimethicone |
| Swiss Navy Silicone Based with Vitamin E | Silicone | Cyclopentasiloxane, Dimethicone, Tocopheryl Acetate (Vitamin E) |

| | | |
|---|---|---|
| Trojan Lubricants Chain Reaction | Silicone | Dimethicone, Dimethiconol, Sensate |
| Wet Platinum | Silicone | Dimethicone, Cyclopentasiloxane, Dimethiconol, Phenyl Trimethicone |
| Wet Water-based | Hybrid | Propylene Glycol, Water, Dimethicone, Cyclopentasiloxane, Hydroxyethylcellulose, PEG/PPG, Caprylhydroxamic Acid, 1,2-Hexanediol, Propanediol, Sodium Polyacrylate, Trideceth-6 |
| Sliquid Silk | Hybrid | Purified Water, Plant Cellulose (from Cotton), Isopropyl Palmitate, Polysorbate 20, Dimethicone, Emollient Ester, Potassium Sorbate, Citric Acid |
| Sliquid Organics Silk | Hybrid | Organic Aloe Barbadensis Leaf Juice, Organic Agar Agar, Organic Guar Gum, Dimethicone, Isopropyl Palmitate, Polysorbate 20, Natural Tacopherois (Vitamin E), Organic Hibiscus Extract, Organic Flax Extract, Organic Green Tea Extract, Organic Sunflower Seed Extract, Potassium Sorbate, Citric Acid |
| BabyDance Fertility Lubricant | Water | Purified Water, Cetyl Hydroxyethylcellulose, Hypromellose, Carbomer Homopolymer Type B, Sodium Phosphate, Potassium Phosphate, Sodium Chloride, Raspberry-derived Xylose, Sodium Hydroxide, Phenethyl Alcohol, Caprylyl Glycol, Salvia Sclarea (Clary Sage) |

| | | |
|---|---|---|
| Conceive + Fertility Lubricant | Water | Water, Hypromellose, Sodium Phosphate, Sodium Dihydrogen Phosphate, Potassium Chloride, Sodium Chloride, Magnesium Chloride, Calcium Chloride, Glycerol, Methylparaben |
